# Diploid donor-specific assembly enhances somatic structural variant detection in cancer genomes

**DOI:** 10.1101/2025.10.28.685155

**Authors:** Yuwei Zhang, Han Qu, Qian Qin, Heng Li, Peter J. Park

**Affiliations:** Department of Biomedical Informatics, Harvard Medical School, Boston, MA 02115, United States; Department of Data Science, Dana-Farber Cancer Institute, Boston, MA 02215, United States; Division of Rheumatology, Inflammation, and Immunity, Brigham and Women’s Hospital, Boston, MA 02467, United States; Broad Institute of MIT and Harvard, Cambridge, MA 02142, United States

**Keywords:** Human Reference Genome, Somatic Alterations, Long-read Sequencing, Genome Assembly, Satellite Repeats

## Abstract

Somatic structural variants (SVs) play a crucial role in tumor development and evolution, yet their identification remains challenging, partly due to limitations in current reference genomes. We present a systematic evaluation of diploid donor-specific assemblies (DSAs)—generated based on hybrid long-read sequencing data—as the reference genome for detecting somatic SVs. We analyzed six tumor-normal cell line pairs, using the EchoSV tool we developed to consolidate haplotype-based SVs into a single DSA-based set and to compare SVs across reference genomes. Across Illumina, PacBio HiFi, and Oxford Nanopore Technology (ONT) data, DSA-based analysis improved read-mapping quality, identified over 20% additional SVs compared to GRCh38 and CHM13, and reduced germline artifacts. Most DSA-specific SVs were isolated deletions or insertions in repetitive elements, especially in satellite regions. By tracking sequence-context differences, we identified scenarios in which DSAs enabled detection of SVs missed on GRCh38/CHM13, and confirmed their functional impact with RNA-seq. These results highlight the value of integrating diploid DSAs into somatic SV analysis.

## Main

Somatic structural variants (SVs) are a pervasive hallmark of human cancer genomes, driving tumorigenesis (1; 2; 3), shaping clonal evolution (4; 5), and modulating therapeutic response (6; 7). For example, in prostate cancer a 3 Mb deletion on chromosome 21 juxtaposes the TMPRSS2 promoter with the ERG coding sequence, producing a TMPRSS2-ERG fusion that aberrantly overexpresses ERG and promotes oncogenesis (8). Similarly, in liver cancer, tandem duplication of the TERT locus leads to multiple copies of the full-length TERT gene, driving telomerase reactivation and cellular immortalization (3). Despite their biological importance, accurate detection of somatic SVs remains challenging owing to intra-tumor heterogeneity (9), extensive repetitive sequence context (10; 11), and mapping ambiguities when aligning to conventional reference genomes (12; 13).

SV identification methods fall into two categories: alignment-based and assembly-based (14). Although *de novo* assembly can resolve rearrangements without reference bias in theory, widespread genome aneuploidy combined with tumor heterogeneity and limited tumor purity render assembly-based approaches impractical for most cancer genomes. Consequently, most cancer genomics studies employ alignment-based calling, where reads are first mapped to a reference genome and then alignments are used to identify variants. In this paradigm, the reference genome is critical, as a genome with unresolved regions or allelic bias can introduce mapping artifacts that mask true SVs or generate false positives (15; 16; 17).

Although GRCh38 has been a dominant reference genome used in the past decade, it was assembled from multiple anonymous donors primarily using Sanger sequencing and still contains over 150 Mb of gaps, mostly in telomeric, centromeric, and long repetitive regions as well as chromosome Y (18). But, with the improved long-read sequencing technologies, notably Pacific Biosciences (PacBio) and Oxford Nanopore Technologies (ONT), near-complete *de novo* assembly has become possible (19). In 2022, the Telomere-to-Telomere (T2T) consortium released CHM13, the first gapless human genome, assembled from a hydatidiform mole using a rich collection of long-read sequencing (15). The authors demonstrated that CHM13 improves germline variant discovery (SNVs, indels, SVs) by resolving previously intractable regions and reducing mapping artifacts. Specifically, CHM13 substantially eliminates spurious homozygous variants in non-African individuals that arise from allelic bias in GRCh38 (roughly 70% of GRCh38 sequences come from a single African-American donor and is enriched for rare and private variants) and exposes variants in centromeric and sub-telomeric regions that are inaccessible in GRCh38. CHM13 has also been shown to enhance somatic SV detection over GRCh38 in the COLO829 tumor-normal-matched sample by a false-positive reduction (17).

Taking a step further, it has now become possible to generate near-T2T-quality diploid assemblies of individual genomes, often termed donor-specific assemblies (DSAs) (20; 21). These *de novo* haplotype-resolved assemblies capture both parental haplotypes and all germline variants, offering a more faithful representation of an individual genome. A previous attempt at using a personalized genome assembly to the normal HCC1395BL cell line reported a >13% increase in somatic SV discovery relative to GRCh38 (22); however, that assembly was haploid (similar to CHM13), where collapsed haplotypes obscure allelic variation.

In this paper, we demonstrate that diploid DSAs enable improved somatic SV discovery compared to GRCh38 and CHM13 across six tumor cell lines with matched normals (Figure 1a). For each pair, we generated a diploid DSA by hybrid assembly of PacBio HiFi and standard ONT reads from the matched normal. Tumor and normal reads were then mapped in a haplotype-specific manner, and somatic SVs were identified separately on each DSA haplotype to allow a head-to-head comparison with the haploid references GRCh38 and CHM13. We developed EchoSV, a tool that captures how SVs “echo” across different references. EchoSV consolidates the two haplotype-based call sets into a single DSA-based set and enables direct comparison with GRCh38- and CHM13-based call sets (Figure 1b; Methods). For each cancer genome, we quantified the somatic SVs revealed only by DSA and traced their origins to sequence-context differences between DSA and conventional references.

**Figure 1.**
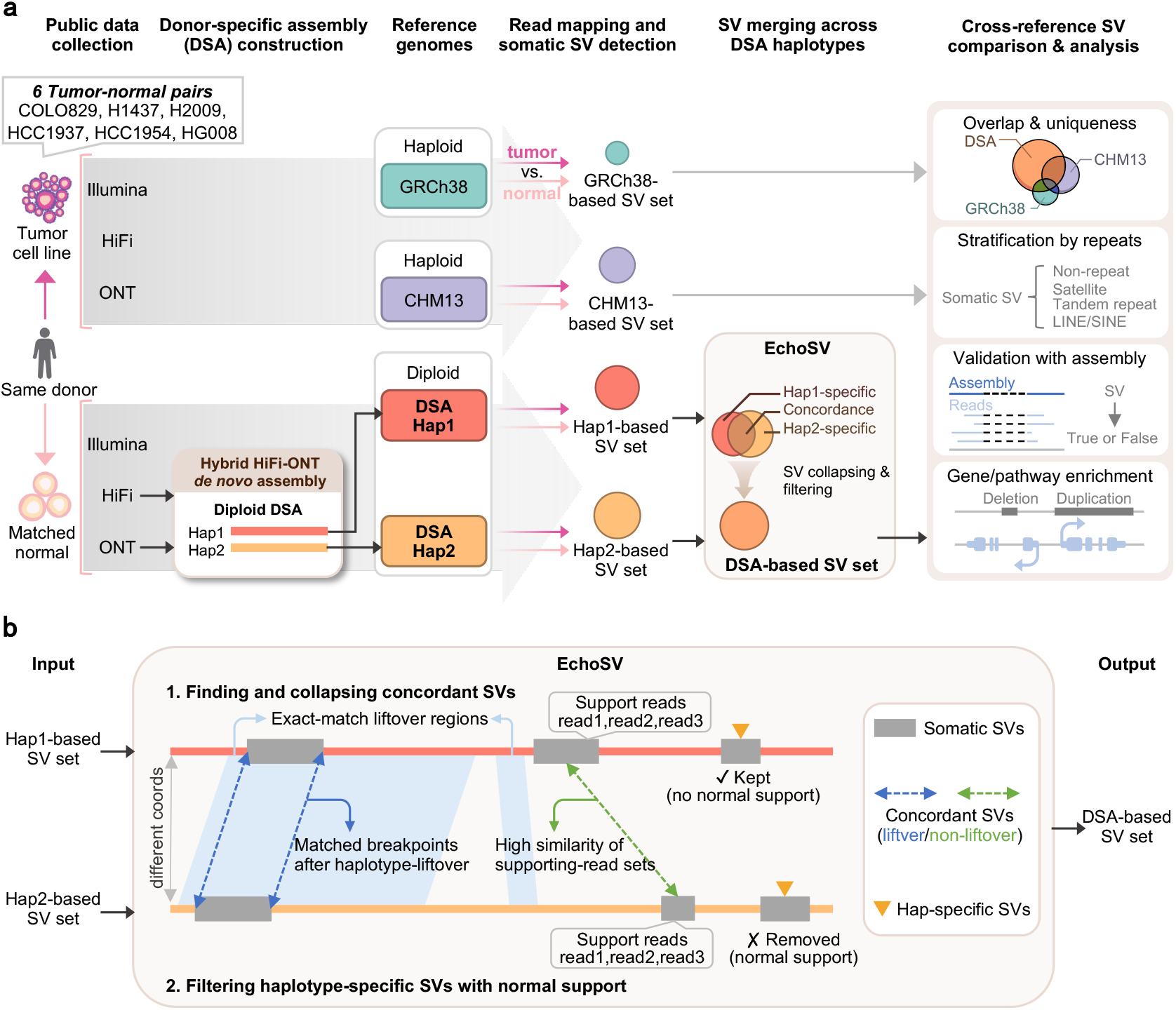
Overview of study design and EchoSV workflow for diploid DSA-based somatic SV analysis. **a**. Experimental design. We analyzed six donor-matched tumor-normal pairs spanning melanoma, lung, breast, and pancreatic cancer. Publicly available Illumina, PacBio HiFi, and ONT reads were collected for each sample. HiFi and ONT reads from each normal sample were assembled into a diploid DSA containing two haplotypes (Hap1 and Hap2). Tumor and matched normal reads were mapped separately to GRCh38, CHM13, Hap1 and Hap2 of DSA, followed by haplotype-specific somatic SV calling. EchoSV is proposed to capture how SVs “echo” across different references. The tool consolidates the two haplotype-based call sets into a unified DSA-based SV set. Downstream analyses compute SV overlap and uniqueness among GRCh38, CHM13, and DSA, annotate shared and reference-exclusive SVs with repeat elements, validate against DSA and tumor assemblies, and intersect genes. **b**. EchoSV workflow. To identify concordant SVs between haplotypes with different coordinate systems, EchoSV first performs haplotype liftover. SVs whose breakpoints align after liftover are classified as concordant. In parallel, EchoSV compares supporting reads across multi-platform sequencing data, and SVs with highly similar supporting read sets are also considered concordant—complementing cases where liftover fails. Concordant SVs are collapsed and reported once in the final DSA-based SV set. SVs present on only one haplotype are subjected to an additional filter to remove germline artifacts before being added to the final DSA-based SV set. EchoSV can be generalized to compare SV call sets across any number of reference genomes.

Compared to GRCh38 and CHM13, diploid DSAs allowed us to detect 45% (median; range: 20%-120%) more somatic SVs across the six tumor-normal pairs. Most DSA-specific events were isolated deletions and insertions within satellite and tandem repeat regions. Below, we detail the quality of the diploid assemblies we constructed, compare read mapping quality across references, evaluate SV detected using multiple algorithms, and characterize the nature and causes of DSA-specific SVs. Our results show that DSAs enable more accurate somatic SV detection and provide practical guidelines for integrating DSA into routine cancer genomics.

## Results

### Construction of near-T2T DSAs for six normal samples

We generated high-quality haplotype-resolved donor-specific assemblies (DSAs) for six human normal samples, including five normal cell lines (COLO829BL, BL1437, BL2009, HCC1937BL, HCC1954BL) and one normal pancreatic tissue sample (HG008-N-P). Hereafter, we refer to these DSAs as COLO829, H1437, H2009, HCC1937, HCC1954, and HG008. Assemblies were constructed with hifiasm (23) using hybrid PacBio HiFi and ONT (Methods). Assemblies for two additional samples—HCC1395BL and HG008-N-D—were suboptimal due to low coverage data and likely sample contamination, respectively, and were excluded from further analysis (Supplementary Table S1).

Despite moderate sequencing depth (median 70x PacBio HiFi and 40x ONT; Supplementary Table S2), all six DSAs achieved contiguity comparable to GRCh38 (version p14, without ALT sequences) and CHM13v2.0 (Table 1). Contig N50 values ranged from 72.3 to 147.6 Mb, better than 52.4 Mb of GRCh38, while scaffold N50 values approached those of GRCh38 and CHM13, despite being derived from more complex diploid genomes. COLO829 exhibited the highest contiguity (scaffold N50: 147.6/134.5 Mb), enabled by deep ONT coverage (140x; the second highest was 50x for H1437). These DSAs substantially outperformed the previous haplotype-collapsed personalized genome assembly HCC1395BL_v1.0 (scaffold N50: 70.0 Mb with 1,645 scaffolds) (22). Although not completely gapless, DSAs contained N content an order of magnitude lower than GRCh38.

**Table 1.** Summary of assembly statistics for GRCh38, CHM13 and six diploid DSAs.

| Assembly | Haplotype | Assembly length (Gb) | N50 (Mb) |  | Ns<sup>a</sup> (Mb) | T2T count |  | QV<sup>a</sup> | Completeness<sup>a</sup> (BUSCO) |
| --- | --- | --- | --- | --- | --- | --- | --- | --- | --- |
|  |  |  | Contig | Scaffold<sup>a</sup> |  | Contig | Scaffold<sup>a</sup> |  |  |
| GRCh38<sup>b</sup> | Haploid | 3.10 | 52.4 | 145.1 | 165.0 | 0 | 0 | NA<sup>c</sup> | 97.9%<sup>d</sup> |
| CHM13 | Haploid | 3.12 | 150.6 | 150.6 | 0 | 24 | 24 | 73.9<sup>e</sup> | 97.9% |
| **COLO829** | Hap1 | 2.95 | 147.6 | 147.6 | 8.1 | 9 | 11 | 58.5 | 98.2% |
|  | Hap2 | 3.05 | 98.2 | 134.5 | 29.9 | 3 | 10 | 59.6 | 93.1% |
| **H1437** | Hap1 | 3.05 | 113.3 | 113.3 | 0.2 | 9 | 9 | 62.2 | 99.1% |
|  | Hap2 | 2.90 | 97.2 | 108.1 | 8.3 | 2 | 9 | 59.8 | 98.6% |
| **H2009** | Hap1 | 3.02 | 93.2 | 103.8 | 5.1 | 3 | 4 | 63.6 | 99.1% |
|  | Hap2 | 3.05 | 92.3 | 103.5 | 2.7 | 4 | 5 | 63.7 | 98.1% |
| **HCC1937** | Hap1 | 2.97 | 102.7 | 112.0 | 0.5 | 7 | 9 | 62.7 | 97.8% |
|  | Hap2 | 2.97 | 99.2 | 111.1 | 10.8 | 6 | 9 | 62.3 | 98.2% |
| **HCC1954** | Hap1 | 3.03 | 110.8 | 127.7 | 6.8 | 3 | 5 | 61.0 | 99.1% |
|  | Hap2 | 3.02 | 89.9 | 127.4 | 6.3 | 3 | 5 | 62.5 | 99.1% |
| **HG008** | Hap1 | 3.02 | 89.6 | 101.4 | 12.2 | 3 | 3 | 62.0 | 98.6% |
|  | Hap2 | 3.02 | 72.3 | 92.2 | 10.6 | 0 | 1 | 61.9 | 98.8% |

<sup>a</sup> Scaffold statistics (N50, T2T count) include both contigs and scaffolds; Ns: the number of unknown nucleotide bases (assembly gaps); QV (quality value): the Phred-scaled consensus base accuracy evaluated by Merqury; Completeness (BUSCO): the percentage of universal single-copy orthologs found intact, evaluated by compleasm
<sup>b</sup> Version p14 without the ALT sequences
<sup>c</sup> NA: not available
<sup>d</sup> The 97.9% value is based on GRCh38.p14 with ALT contigs
<sup>e</sup> The 73.9 value is based on CHM13v1.1 with Merqury

Our DSAs achieved a median of 14 T2T contigs and scaffolds across samples, comparable to the latest Human Pangenome Reference Consortium (HPRC) release, which reported a median of 18 T2T contigs and scaffolds despite leveraging not only HiFi and ONT but also ultra-long ONT reads (>100 kb) and Strand-seq (24).

The six DSAs also exhibited high base-level accuracy, with quality values (QV) ranging from 58.5 to 63.7, corresponding to an error rate of 0.4-1.4 per million bases (25). Completeness assessments using BUSCO (26) revealed extremely high gene content (>98.2%), exceeding that of GRCh38 and CHM13 (97.9%), and consistent with near-reference completeness despite being haplotype-resolved.

Together, these results demonstrate that integrating moderate-coverage HiFi and ONT data can produce near-reference-quality, haplotype-specific assemblies suitable for use as personalized genome references.

### Sequence comparisons of DSAs with GRCh38 and CHM13

We compared each DSA with GRCh38 and CHM13 by aligning both haplotypes using minimap2 in assembly-to-assembly alignment mode (28). To characterize how DSAs diverge from standard references, we classified reference regions into four categories based on alignment coverage: uniquely mappable (aligned by exactly one alignment), unmappable (no alignment), duplicated (two alignments), and highly duplicated (>2 alignments) (Supplementary Figure S1). Across the six DSAs, an average of 92.0% (±0.6%) of GRCh38 and 94.8% (±1.1%) of CHM13 were uniquely mappable by at least one DSA haplotype, reflecting a high degree of shared sequence. Divergent regions were largely confined to donor-specific sequences and assembly-challenging loci. Taking COLO829-to-CHM13 alignment as an example (Figure 2a), the X-q arm is unmappable by Hap1, consistent with the male donor. Smaller unmappable (grey) or duplicated (yellow) regions likely represent germline deletions and duplications on CHM13, whereas highly duplicated regions (blue) were enriched in pericentromeric repeats and likely reflect assembly challenges and mapping artifacts. We defined differential reference regions (DRRs) on CHM13 as segments not uniquely mappable by either COLO829 haplotype and found these DRRs co-localize with telomeres, centromeres, acrocentric p-arms, and chromosome Y, all known challenges for *de novo* assembly (29). In the comparison with COLO829, GRCh38 exhibited a notably larger DRR fraction than CHM13 (7.4% vs. 4.1%), driven largely by unresolved Ns in GRCh38 (Supplementary Figure S2).

**Figure 2.**
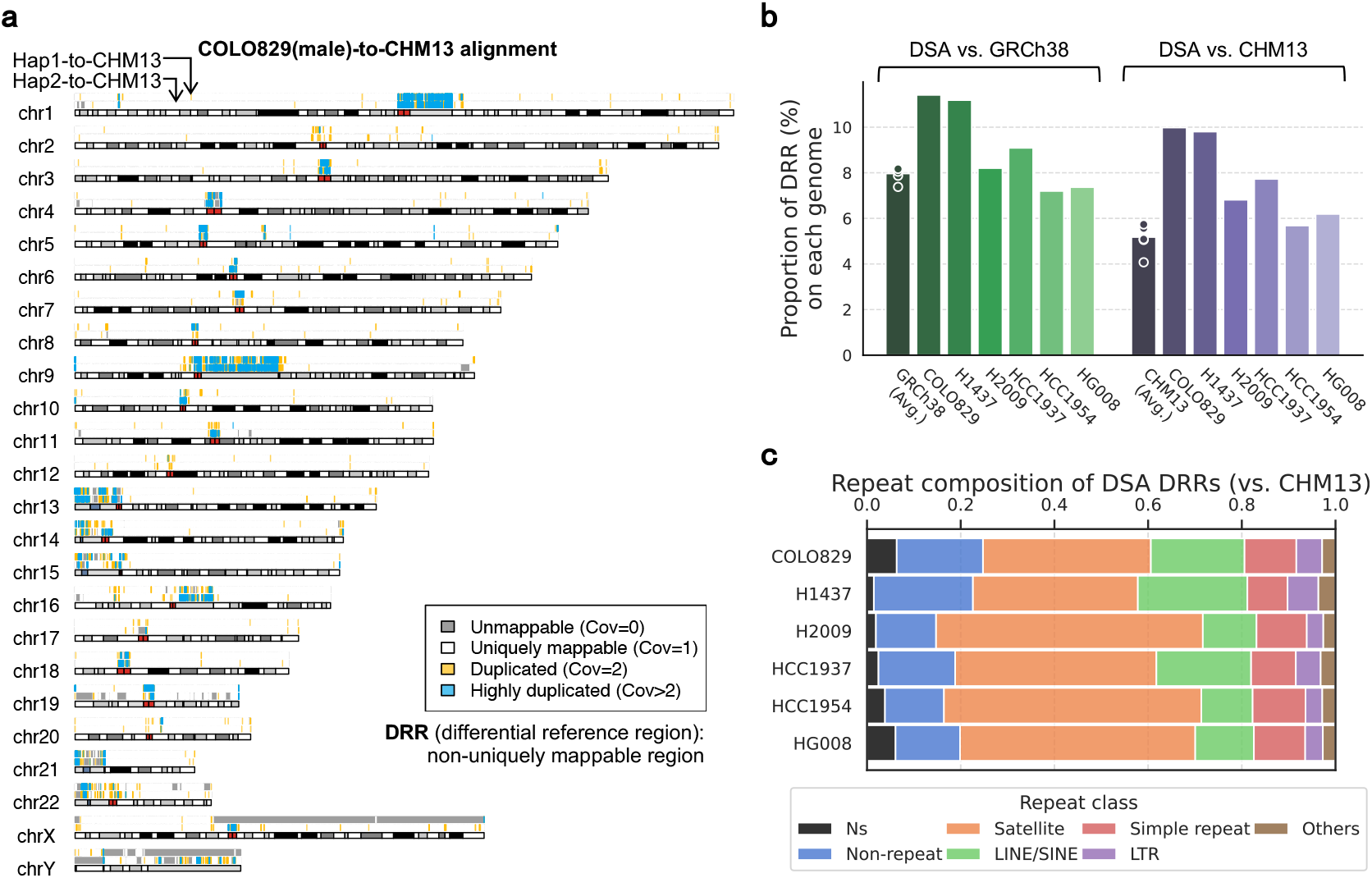
Comparison of DSA sequences with GRCh38 and CHM13. **a**. Assembly-to-assembly alignments of both haplotypes of the COLO829 DSA (male donor) against CHM13 using minimap2, visualized with KaryoploteR (27). Four categories of regions on CHM13, defined by alignment coverage, are shown in different colors. **b**. Proportion of differential reference regions (DRRs) in pairwise DSA-GRCh38 (green) and DSA-CHM13 (purple) comparisons. For “GRCh38” and “CHM13”, bar heights denote the average DRR proportion across six DSA comparisons; dots represent values from individual comparison. **c**. Repeat composition of DSA DRRs relative to CHM13, annotated by RepeatMasker. Stacked bars show the fraction of each repeat class per DSA; repeat classes are ordered by average composition across samples except “Ns” and “Non-repeat”. Abbreviations: Ns, assembly gaps; LINE, long interspersed nuclear element; SINE, short interspersed nuclear element; LTR, long terminal repeat.

We next generalized the definition of DRRs to all pairwise sequence comparisons. Whereas uniquely mappable regions are symmetric between two genomes, DRRs are inherently asymmetric: in the COLO829-to-CHM13 comparison, DRRs on CHM13 are segments not uniquely mappable by both COLO829 haplotypes, while DRRs on COLO829 are segments not uniquely mappable by CHM13. Thus, DRRs capture both novel and duplicated sequences specific to each genome that cannot be lifted over. Across all six DSAs, GRCh38 exhibited a higher mean DRR proportion than CHM13 (8.0% versus 5.2%; Figure 2b). Similarly, DSAs showed a larger proportion of DRRs against GRCh38 than against CHM13 (9.1% vs. 7.7%, or 545 Mb vs. 463 Mb on average across six samples), consistent with CHM13 providing a more complete and less biased representation of the human genome (15).

To examine the sequence composition of DSA DRRs (344-600 Mb per DSA compared to CHM13), we annotated them using RepeatMasker (30) (Figure 2c). Satellite repeats constitute the largest fraction (35-57%, mean 46%), followed by LINEs/SINEs (11-23%, mean 16%) and simple repeats (9-11%, mean 10%). Together with non-repetitive sequence (13-21%, mean 16%), these categories accounted for over 88% of all DRR bases. The remainder consisted of assembly gaps and other repeat elements, such as LTRs. Although repeat-rich DRRs pose challenges for read mapping and variant calling, they harbor a wealth of donor-specific sequences that can enable the discovery of novel SVs and their functional interpretation.

### DSA enhances SV-related read mapping quality metrics

We assessed the impact of DSAs on read mapping relative to GRCh38 and CHM13. Sequencing reads from each tumor–normal pair were aligned to four references—GRCh38, CHM13, DSA hap1 and DSA Hap2—independently,using BWA-MEM (31) for Illumina and minimap2 (28) for HiFi and ONT. The haplotype-specific mapping strategy with DSAs avoids misalignment caused by homologous sequences (the two haplotypes share 99.5% identity) (32). For metric calculations, alignments from both DSA haplotypes were combined.

We evaluated three metrics relevant to the subsequent SV calling’s sensitivity and precision. Q10 mappability was defined as the fraction of reads (or read pairs for Illumina) with MAPQ*≥*10; higher values indicate more high-confidence alignments and likely higher SV calling sensitivity. Compared with GRCh38 and CHM13, DSAs showed a clear increase in Q10 mappability for long reads (Figure 3a, right two columns). In contrast, gains for short reads were minimal, suggesting that DSAs mainly improve alignment within repetitive regions.

**Figure 3.**
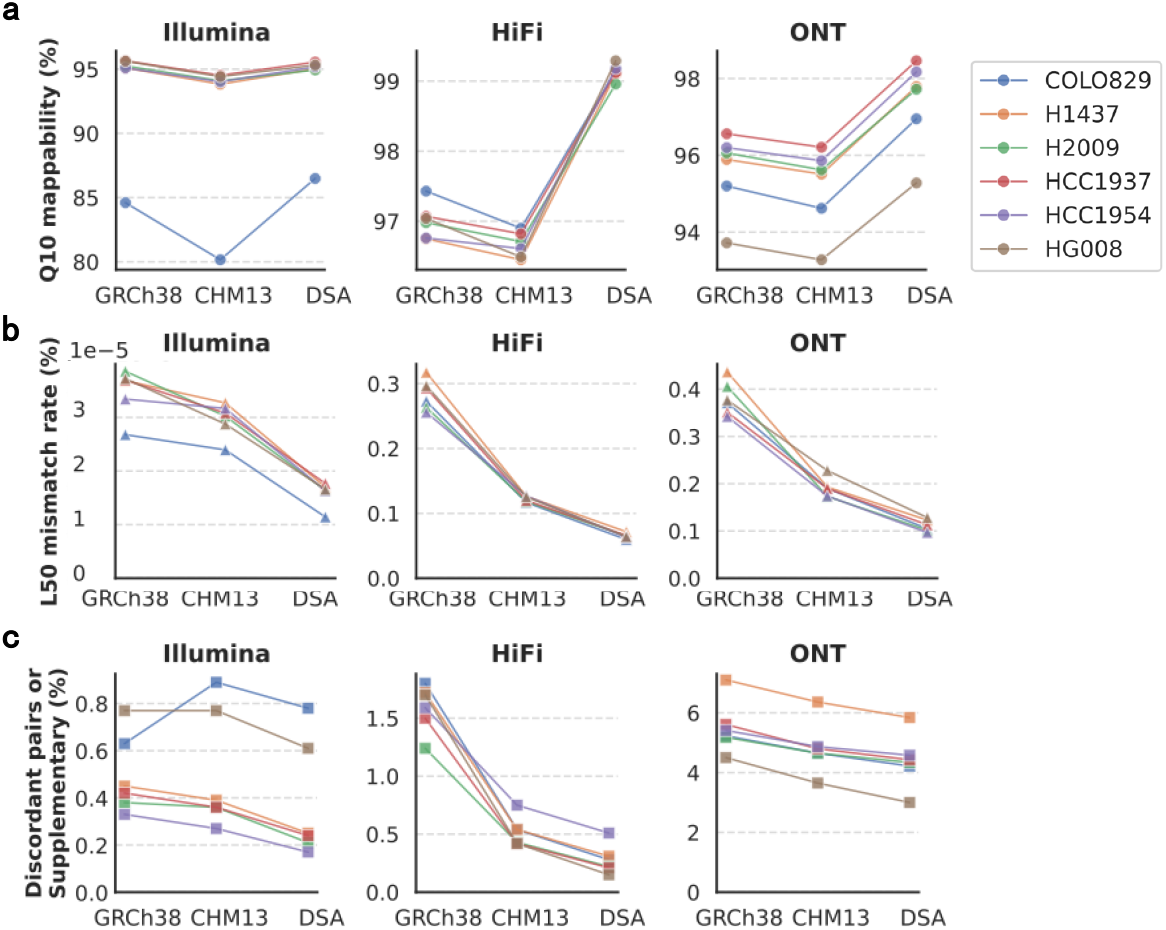
Comparison of tumor reads mapping performance on GRCh38, CHM13 and DSAs, using SV-related metrics. **a**. Q10 mappability (fraction of reads or read pairs with MAPQ *≥*10), **b**. L50 mismatch rate (fraction of bases aligned with insertions, deletions, or mismatches *≥*50 bp), and **c**. inter-alignment SV signal rate (fraction of discordant read pairs and split reads with MAPQ *≥*10). Colors denote individual tumor samples as indicated in the legend of **a**.

We defined the L50 mismatch rate as the fraction of bases within MAPQ*≥*10 alignments where the CIGAR string records an insertion, deletion, or mismatch of *≥*50 bp (intra-alignment SV signal rate). Lower values indicate fewer insertion- and deletion-like signals and reduced alignment artifacts. For both short and long reads, alignments to DSAs showed substantially lower L50 mismatch rates than those to GRCh38 and CHM13 (Figure 3b), underscoring their potential to improve INDEL (insertions and deletions of *≥*50 bp) calling precision across all sequencing platforms.

Detection of inversions, duplications, and translocations relies on split-read alignments and, for Illumina, discordant read pairs (33; 34; 35). We quantified these signals as the fraction of MAPQ*≥*10 reads that were either split at least or classified as discordant pairs (inter-alignment SV signal rate). For both short and long reads, DSAs showed lower fractions relative to GRCh38 or CHM13 (with the only exception of short-read mapping in the COLO829 sample; Figure 3c), indicating reduced background noise and improved precision for detecting SVs beyond simple INDELs.

### More SVs are identified using DSA than using GRCh38 or CHM13

To obtain high-confidence SVs, we applied a multi-tool, multi-platform pipeline to each tumor-normal pair on every reference haplotype (Figure 4a). Fifteen alignment-based calling strategies—each a combination of a caller with a short- or long-read platform—were run on GRCh38, CHM13, DSA Hap1, and DSA Hap2 (Methods). A variant entered the high-confidence set only if it was supported by *≥*2 sequencing platforms (of 3) and *≥*4 callers (of 15). The number of high-confidence somatic SVs on GRCh38 exceeded that of previous ensemble set using the same thresholds on the same sample (12) (Supplementary Table S3), owing to our expanded set of callers and updated caller versions.

**Figure 4.**
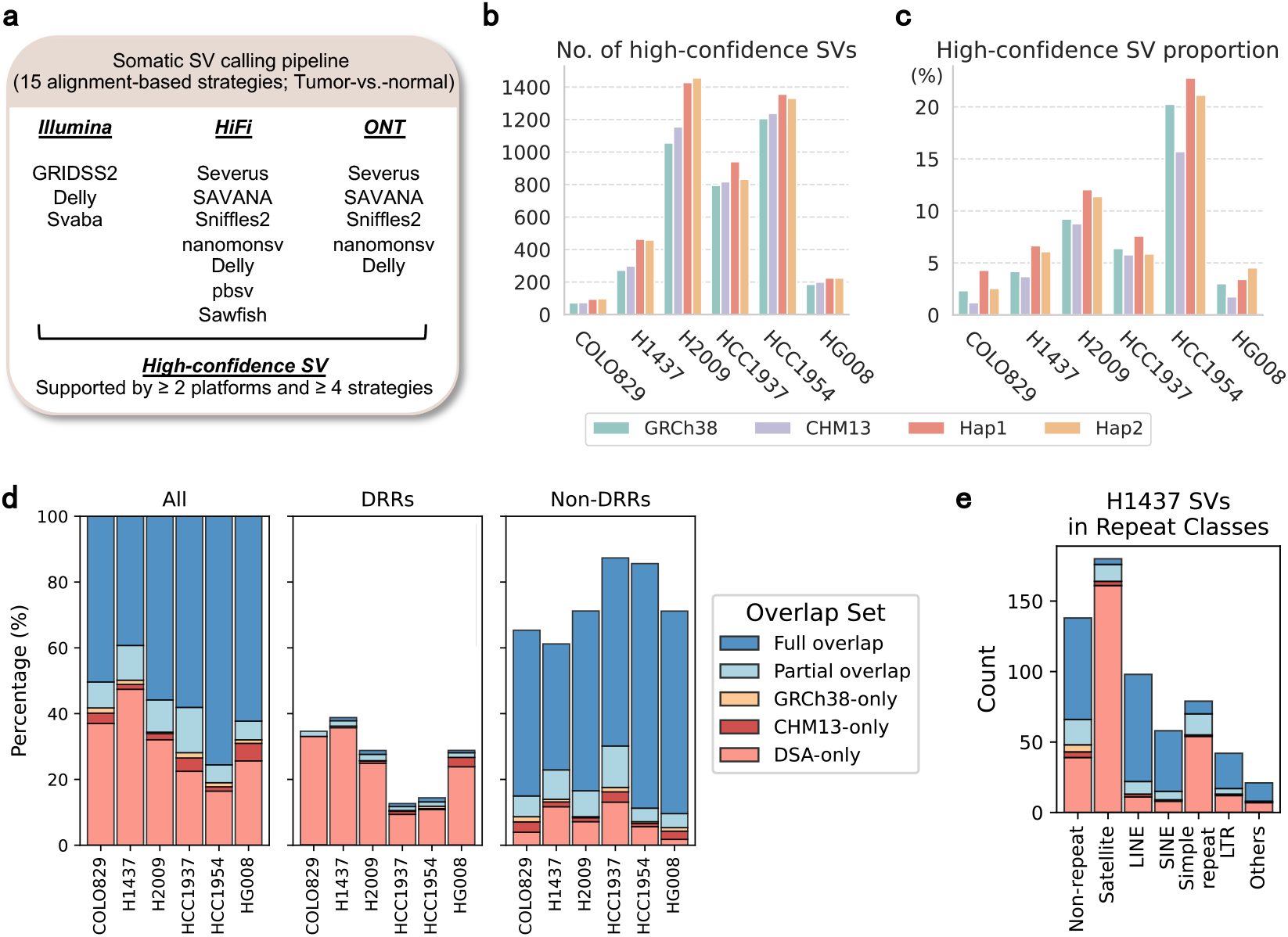
DSA enhances somatic SV discovery. **a**. Multi-tool, multi-platform somatic SV calling pipeline applied to each tumor-normal pair and reference. Fifteen alignment-based calling strategies spanning three sequencing platforms and nine callers were used. High-confidence SVs were derived from the union of the call sets meeting support criteria. **b**,**c. b**. Counts and **c**. proportions of high-confidence somatic SVs detected on GRCh38, CHM13, DSA Hap1 and DSA Hap2. **d**. Fraction of SVs per cell line classified as full overlap (present in all GRCh38, CHM13, and DSA), partial overlap (present in two references), or reference-exclusive, shown for the whole genome, DRRs, and non-DRR regions. **e**. SV counts in the cell line with the largest DSA-only fraction (H1437), stratified by RepeatMasker class.

Both Hap1 and Hap2 yielded 2-70% (mean 27%) more high-confidence calls than GRCh38 or CHM13 across all samples (Figure 4b). Furthermore, the fraction of high-confidence SVs among the union of all fifteen call sets was also higher on DSA haplotypes (Figure 4c), except for HCC1937 Hap2. These results indicate that, as expected from improved mapping quality described above, DSAs better account for individual-specific variations and difficult-to-align regions while reducing caller- and platform-specific noise.

Next, we derived DSA-based SV sets by consolidating haplotype-based calls to enable a more proper comparison with GRCh38 and CHM13. This was achieved with our new tool EchoSV (Figure 1b; Methods), which merges concordant SVs detected on both haplotypes through both liftover and read-support similarity, while retaining haplotype-exclusive SVs with no support in the matched normal. After consolidation, DSA-based high-confidence sets contained, on average, 58% more somatic SVs than GRCh38 and 49% more than CHM13 (Supplementary Table S4). By SV type, DSAs uncovered 102% more deletions, 33% more insertions/duplications, and 36% more translocations, whereas inversion counts were similar across references (Supplementary Figure S3).

With EchoSV, we classified high-confidence SVs across GRCh38, CHM13, and DSA into five categories: full overlap (present in all three references), partial overlap (shared by exactly two), and three reference-exclusive classes (Figure 4d, left panel). Across samples, DSA-only calls comprised 16-47% of all SVs, outnumbering GRCh38-only plus CHM13-only calls by eight-fold (Supplementary Table S5). Among SVs shared by two references, 92% involved DSAs (58% DSA+CHM13, 34% DSA+GRCh38), while only 8% were shared between CHM13 and GRCh38, further underscoring DSA’s superior sensitivity for detecting somatic SVs missed by standard references. Not surprisingly, these DSA-only SVs were more enriched within DRRs (Figure 4d, middle and right panels). Despite variation in repeat composition across samples, DSA-only SVs were consistently and highly represented in satellite repeats (Figure 4e; Supplementary Figure S4).

### DSA provides an abundance of novel somatic SVs in satellites and tandem repeats

We next examined reference-exclusive high-confidence SVs using assembly-based validation (Figure 5a; Methods). In addition to the constructed DSAs, we assembled each tumor genome with PacBio HiFi reads using hifiasm (36) and aligned the resulting tumor assemblies to the reference. As a stringent criterion, a reference-exclusive SV was accepted as a true positive only if reproduced in the tumor assembly alignment. For variants absent from the tumor assembly, we further assessed whether they were present in DSA alignments to the reference. Two error modes were identified: germline artifacts (SVs present in the DSA alignment) and mapping artifacts (non-insertion SVs co-localizing with breakpoints of large germline insertions, reflecting mis-mapping). Variants not validated as either true or false positives were considered unknown.

**Figure 5.**
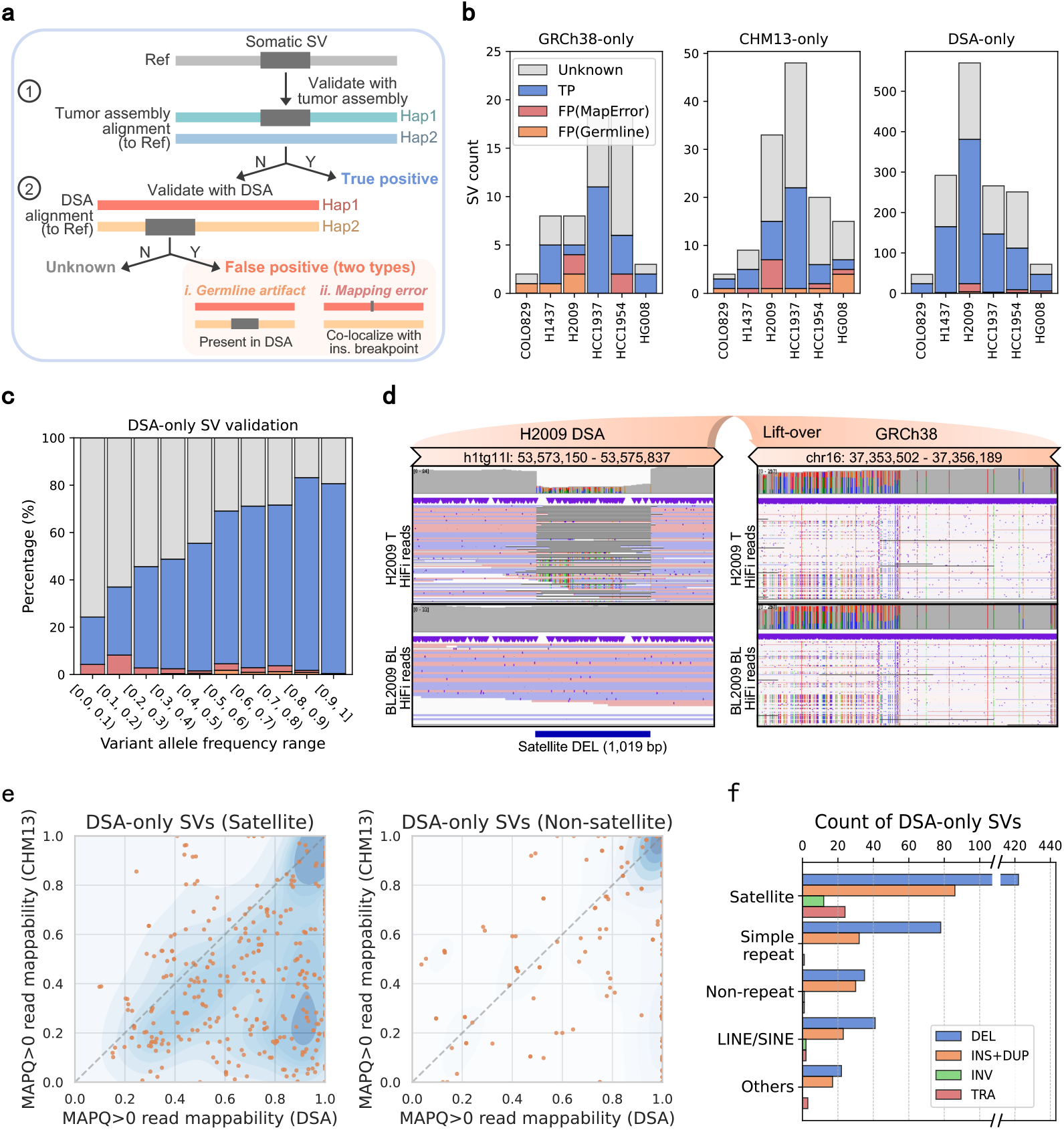
Examination of reference-exclusive SVs using DSA and tumor assemblies. **a**. Assembly-based validation strategy. Each somatic SV was classified by validation status based on tumor and DSA alignments: true positives (reproduced in tumor assemblies), false positives (germline if reproduced in DSAs, or mapping artifacts if co-localizing with germline insertion breakpoints thus partially reproduced in DSAs), and unknown events (not validated as either true or false positives). **b**. Stacked bar plots showing counts of reference-exclusive SVs by validation status. **c**. Percentage of DSA-only SVs with different validation status, stratified by 0.1 variant allele frequency (VAF) bins. Legend is the same as in **b**. **d**. IGV screenshot of a 1,019 bp somatic deletion in a centromeric satellite repeat (H2009, h1tg11l:53,574,276–53,575,295). The deletion was confirmed in the tumor assembly and detected only on DSA, but missed on GRCh38 due to poor mappability. Reads are colored pink (forward strand) and purple (reverse strand); white reads indicate MAPQ=0. **e**. Fraction of reads with MAPQ>0 within 500 bp window of validated DSA-only breakpoints, comparing DSA versus CHM13 alignment quality for SVs in satellite and non-satellite regions. Each dot represents one DSA-only breakpoint. Dot density was estimated using kernel density estimation (KDE), with darker regions indicating higher density. **f**. Count of DSA-only SVs in repeat classes by SV type. Repeat classes are ordered by total SV count. Abbreviations: DEL, deletion; INS, insertion; DUP, duplication; INV, inversion; TRA, translocation.

As expected due to subclonal heterogeneity, the tumor assemblies were highly fragmented, with 700 to 4,900 contigs per sample and N50 values ranging from 26.7 to 72.3 Mb (Supplementary Table S6). Even so, 54% of DSA-only SVs were validated as true positives, compared with only 30% of GRCh38- or CHM13-only SVs (Figure 5b; Supplementary Table S7). This disparity is largely explained by a sharp reduction in false positives. On average, only 3% of DSA-only SVs were identified as germline variants or mapping artifacts—a greater than five-fold reduction over the 19% and 17% for GRCh38-only and CHM13-only SVs, respectively. This low false-positive rate is consistent with recent work(37). The reduction in germline errors was particularly striking: 0.5% for DSA-only SVs versus 10.3% for those unique to GRCh38 and CHM13. The remaining unknown 622 DSA-only SVs (43%) exhibited lower variant allele fractions (VAFs) and similar SV signature profiles to the validated true positives, suggesting they are likely genuine (Supplementary Figure S5). These validation results demonstrate that DSAs provide high sensitivity and precision for somatic SV detection with negligible loss of true positives exclusively on standard references.

DSA also excelled in detecting SVs within satellite repeats, where 66% of 824 DSA-only satellite SVs were validated as true positives. In contrast, only 22% of CHM13-only satellite SVs (typically one per sample) and none of GRCh38-only satellite SVs could be confirmed (Supplementary Figure S6). Because low VAF impairs assembly, we binned DSA-only SVs in 0.1 VAF intervals and measured the associated validation rates. We observed a steady rise in validation rates with increasing VAF (Figure 5c), consistent with expectations and reinforcing the reliability of our assembly-based validation strategy.

To understand why only DSAs reliably capture somatic SVs in satellite repeats, we first examined individual cases. A representative example is a 1,019 bp somatic deletion within a centromeric satellite repeat in H2009 (Figure 5d). On DSA, tumor read alignments display a clear coverage drop and deletion-supporting evidence absent in the matched normal sample. On GRCh38, however, the same reads show low mapping quality (many MAPQ=0 reads in white), fragmented alignments with scattered indels, and dense mismatches, leaving no clear somatic signal across the 3kb flanking region. This observation holds true systematically: we compared the fraction of non-zero MAPQ reads within 500 bp of each validated DSA-only breakpoint, and found that read mappability around satellite SV breakpoints was substantially higher on DSA compared to the corresponding CHM13 coordinates (Figure 5e). For non-satellite DSA-only SVs, mappability was uniformly high on both references (high density on the top-right corner). Importantly, we reproduced the majority of these satellite SVs using an alternative assembler(Verkko(38), with available Pore-C data) and aligner (Winnowmap(39)), confirming the robustness of our findings (Supplementary Figure S7).

These results are consistent with the known biology of satellite DNA, which suppresses homologous recombination (HR) by creating a physical barrier for HR factors and thus favors error-prone non-homologous end joining (NHEJ), making these regions hotspots for somatic SVs (40; 41). Indeed, our results show a prevalence of deletions here (Figure 5f). These regions are typically excluded from analyses using GRCh38 and CHM13 due to excessive mapping artifacts(9; 17). Our results demonstrate that by restoring mappability in these repeat-rich regions, diploid DSAs uncover a large and previously hidden reservoir of somatic SVs in cancer genomes.

### Case studies of DSA-specific SVs

We highlight four scenarios in which reference-context differences underlie DSA-specific SV detection, using representative assembly-validated examples across diverse repeat classes. Figure 6a involves reference contraction, where a standard reference contains a shorter version of a repeat than DSA. The 60 bp homozygous deletion in an (AT)_*n*_ repeat identified on DSA was missed when using GRCh38 and CHM13 because both DSA haplotypes were 30 bp longer (15 AT units) than GRCh38 and CHM13. Consequently, the true 60 bp somatic deletion was misinterpreted as a 30 bp event, falling below the 50 bp size threshold and therefore undetectable. Similar effects were observed in other tandem repeats, including (TA)_*n*_, (TTC)_*n*_ and (ATATATATA_*n*_.

**Figure 6.**
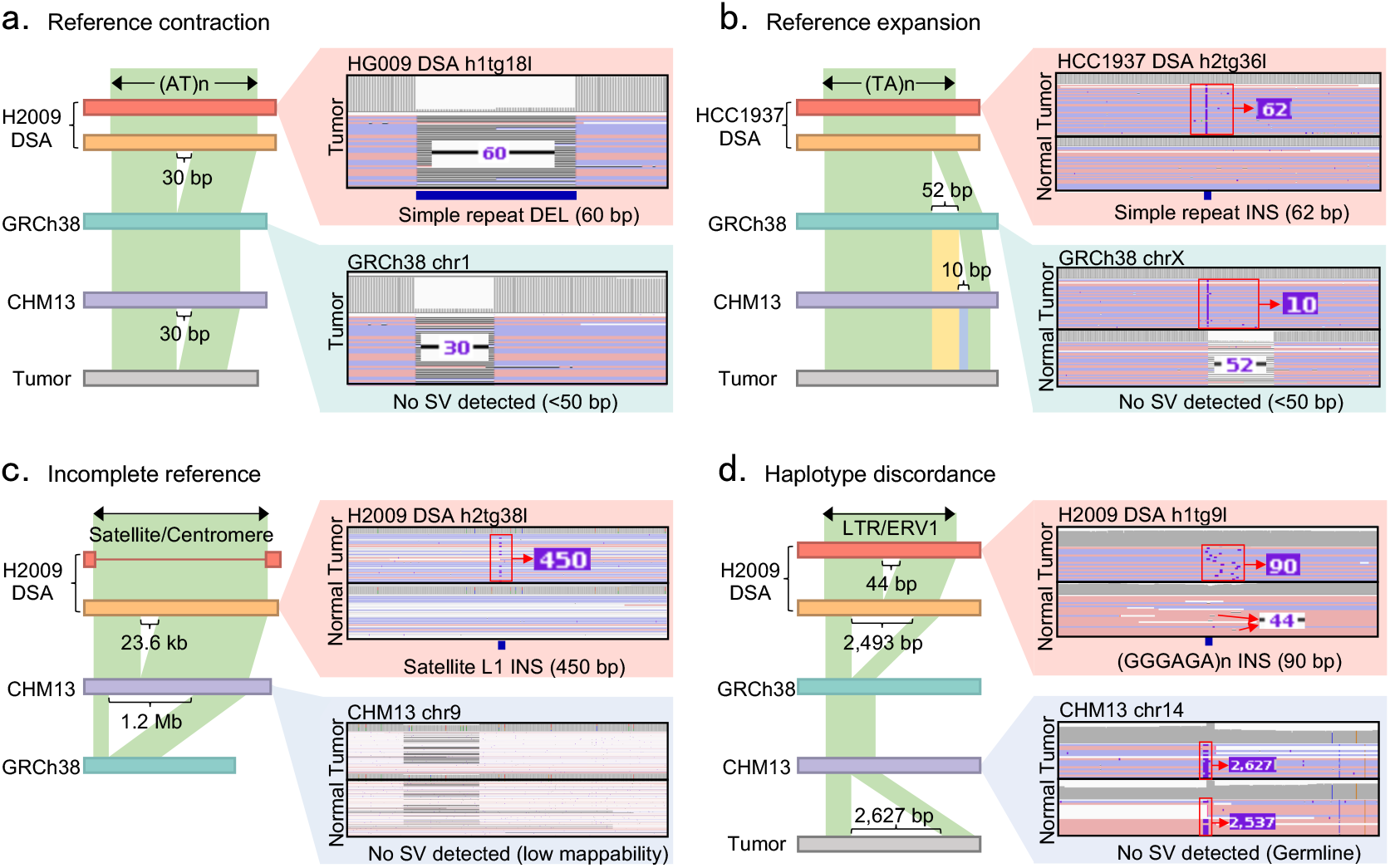
DSA resolves SVs missed due to reference discrepancies. **a. Reference contraction**. A 60 bp deletion in an (AT)_*n*_ repeat on H2009 DSA (h1tg18l:25,829,176–25,829,236) is undetectable on GRCh38 and CHM13 because both references are 30 bp shorter than the donor’s germline. The true 60 bp event is misinterpreted as a 30 bp deletion agains GRCh38 and CHM13, falling below the 50 bp detection threshold. **b. Reference expansion**. A 62 bp insertion on HCC1937 DSA (h2tg36l:13,815,463) is missed on GRCh38 and CHM13, which both contain a longer repeat allele relative to the donor’s germline, masking the somatic event and causing it to be missed. **c. Incomplete reference**. A 450 bp L1 insertion within a centromeric satellite on H2009 DSA (h2tg38l:39,702,182). This satellite spans 3.21 Mb in the DSA but 3.18 Mb in CHM13 and only 0.65 Mb in GRCh38, leading to low mappability (many MAPQ=0 reads) and artifactual deletion signals in both tumor and normal read alignments that obscure the true event. **d. Haplotype discordance**. A 90 bp insertion on H2009 DSA (h1tg9l:11,438,901). Hap1 contains a 44 bp segment absent from Hap2, so the event appears as a 90 bp insertion on Hap1 and a 134 bp insertion on Hap2. GRCh38 and CHM13 are 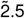 shorter at this locus, preventing confident variant detection due to minor insertion-size-difference between tumor and normal sample. For each case, IGV snapshots of PacBio HiFi tumor (and normal) read alignments are shown on DSA and at the corresponding loci on GRCh38 or CHM13, illustrating how reference-specific context affects SV detection. Reads are colored by strand (pink: forward, purple: reverse), with white indicating ambiguously mapped reads (MAPQ=0).

Conversely, reference expansion—where the standard references contain a longer repeat than the donor’s germline—also masks true SVs (Figure 6b). In one such case, DSA correctly identified a 62 bp homozygous insertion. However, because GRCh38 already contained a 52 bp expansion at this locus relative to the donor, the event was shown as a small 10 bp insertion and therefore missed. Against CHM13, which harbored an expansion matching the 62 bp somatic insertion, the event was completely obscured, resulting in no SV detected.

A third class of SVs missed by GRCh38 and CHM13 arises from incomplete reference sequences, particularly within large satellite repeats(Figure 6c). We identified a 450 bp somatic L1 insertion in a centromeric satellite that was only detectable using DSA. The satellite region spanned 2.67 Mb in DSA, 2.65 Mb in CHM13, and only 1.46 Mb in GRCh38. This reference incompleteness caused reads originating from this region to map ambiguously (most received MAPQ=0) on GRCh38 and CHM13, creating strong artifactual deletion signals in both tumor and normal samples. In contrast, DSA provided unique anchoring sequences that enabled high-quality mapping and confident detection of this true somatic mobile element insertion (also validated using Verkko-generated DSA and Winnowmap alignments; Supplementary Figure S8). Similarly, we found 27 such L1 insertions within satellite regions in the H2009 cell line (all with poly-A tail and target site duplication signals), revealing a prevalent yet previously uncharacterized pattern of somatic retrotransposition. This example again underscores the value of DSAs for variant detection in repeat-rich, polymorphic regions such as satellites.

The final scenario demonstrates how DSA enables SV detection in haplotype-discordant regions (Figure 6d). We identified a 90 bp somatic insertion that was missed by GRCh38 and CHM13 because they both lacked a 2,500 bp segment at this locus. This misrepresented the event on CHM13 as two large, similarly-sized insertions in the tumor (2,627 bp) and normal (2,537 bp) samples, falling below somatic calling thresholds. Furthermore, DSA revealed that this locus was discordant between the donor’s haplotypes: a 44 bp segment present in Hap1 but absent from Hap2 caused the event to manifest as a 90 bp insertion on Hap1 and 134 bp on Hap2. EchoSV correctly recognized these as concordance and collapsed them into a single somatic SV. Notably, 45% of DSA-only SVs occurred in such haplotype-discordant regions, variants that haplotype-collapsed assemblies as GRCh38 and CHM13 would likely miss. This example illustrates the advantage of diploid DSAs and the accuracy of EchoSV in resolving complex, haplotype-specific SVs.

The 60 bp deletion in Figure 6a disrupted an intron of STX12, a SNARE protein linked to cell invasiveness (42). RNA-seq data indicated STX12 downregulation in H2009, and overall, 12% of DSA-only SVs overlapped genes and 44% of these genes were differentially expressed in tumors relative to matched normals (Supplementary Figure S9). Functional enrichment highlighted pathways related to epidermal differentiation and peptide cross-linking, processes associated with oncogenesis. These results suggest the functional insights enabled by novel variants uniquely detected with diploid DSAs.

## Discussion

The algorithms for somatic SV detection have proliferated over the past 15 years. A 2021 study evaluated 7 tools for short-read data (10), while a 2025 study assessed 16 long-read-based pipelines combining long-read platforms with diverse detection methods (12). But their performance is highly variable—substantially more so than for SNV detection—across tools and datasets. Several factors influence algorithm performance, including variant characteristics (type, size, VAF), sequencing data (read length, depth), and various threshold choices (minimum read support, mapping quality) for the trade-off between sensitivity and specificity. In this work, we highlighted another, previously underexplored source of variability: the reliance on haploid reference genomes that differ substantially from an individual’s diploid genome. Without accounting for individual genomic differences, deriving a comprehensive set of true somatic SVs becomes nearly impossible, leaving evaluations of algorithm performance inherently incomplete.

The DSAs we constructed yielded roughly 500 Mb of sequences absent from GRCh38 but specific to the donor’s haplotypes, mostly composed of satellite DNA and other repeat-rich elements. Incorporating these sequences enhanced read mapping by increasing mappability and reducing artifacts, as reads from donor-specific regions are often misaligned on GRCh38 or CHM13, and it facilitated discovery of novel SVs in satellites and otherrepeats (particularly with long-read data; Supplementary Figure S10). Overall, we were able to identify 49% more somatic SVs on average compared to using CHM13 and validated more than half of these DSA-only SVs through generated tumor genome assemblies. Notably, the observed enrichment of SVs in satellite regions is consistent with the previous findings that heterochromatin more frequently undergoes somatic structural rearrangements via NHEJ (40; 41).

Until recently, generating a high-quality diploid genome was technically challenging and prohibitively expensive, as multiple sequencing data types of sufficient quality were required. PacBio HiFi reads offer high accuracy but inadequate read length for resolving highly repetitive regions, whereas ONT provides longer reads but with lower base-calling accuracy. In this paper, we utilized an improved algorithm available in hifiasm (23) to a combination of PacBio and ONT reads to obtain near-T2T-quality DSAs. Recent efforts from our groups suggest that high-quality ONT data alone may now suffice for generating accurate DSAs (43), which would substantially lower the sequencing costs. These developments and our analysis indicate that routine analysis of diploid DSA may be possible in the near future. As more samples are subjected to DSA analysis, we will have more opportunities to elucidate the biological mechanisms underlying DSA-specific SVs and their functional consequences.

## Methods

### Publicly available data of tumor-normal pairs

We analyzed six tumor-normal cell-line pairs—COLO829, H1437, H2009, HCC1937, HCC1954 and HG008—for which sequencing data are available from all three major platforms: Illumina, PacBio HiFi, and standard ONT. Both PacBio HiFi and ONT long reads are required for DSA generation, while each platform independently supports somatic SV calling.

Most data were retrieved from BioProject PRJNA1086849 (12), including Illumina and ONT for COLO829, data from all three platforms for H1437, H2009, HCC1937, and HCC1954. PacBio HiFi data for COLO829 were downloaded from PacBio public repository (44). All sequencing was performed on state-of-the-art instruments—Illumina NovaSeq 6000, PacBio Revio, and ONT PromethION R10 flow-cells—except for the COLO829 ONT dataset, which used R9.4.1 flow-cells.

The HG008 samples were obtained from GIAB (Genome in a Bottle Consortium) (45). HG008-T is a pancreatic ductal adenocarcinoma cell line, with matched normal pancreatic (HG008-N-P) and duodenal (HG008-N-D) tissues from the same donor. We analysed passage-23 HG008-T reads with combined Illumina data from BCM and NYGC, HiFi data from BCM and PacBio, and both standard and ultra-long ONT reads from UCSC. For HG008-N-P, we used HiFi plus standard ONT R10 data; for HG008-N-D, we merged Illumina data from BCM and NTGC, HiFi data from BCM, and standard ONT R10 data. Coverage summaries for all raw datasets are provided in Supplementary Table S2.

Gene expression TPM (transcripts per million) table of protein-coding genes for tumor cell lines was downloaded from the DepMap (Dependency Map, 24Q4) (46). The table is inferred from RNA-seq data using GTEx pipelines, with values log2-transformed and batch-corrected. For normal samples, we downloaded the median TPM of non-diseased tissues from GTEx (v10)(47) and paired these normal tissues to each tumor cell line.

### Donor-specific assembly construction and quality control

PacBio HiFi and ONT reads from each normal sample (COLO829BL, BL1437, BL2009, HCC1937BL, HCC1954BL, HG008-N-P) were assembled with hifiasm (version 0.20.0-r639) (36). HiFi reads served as the primary input, with ONT data integrated as ultra-long reads (`–-ul`) to improve contiguity across challenging genomic regions. The ultra-long read length cutoff `(–-ul-cut`) was tuned per sample: 50 kb for COLO829, 30 kb for H1437, and 10 kb for HCC1937, HCC1954, H2009, and HG008-N-P. The parameter `–-dual-scaf` enabled scaffolding capabilities and `–-telo-m CCCTAA` preserved telomeric repeats.

A comprehensive evaluation was performed with complementary approaches to assess assembly completeness, contiguity, and accuracy. T2T contigs and scaffolds were identified with the HPRC (Human Pangenome Reference Consortium) workflow (CHM13 v2.0 reference)(48). Contiguity was assessed by contig and scaffold N50/N90 values. Base accuracy was estimated by yak (v0.1-r69-dirty)(49) using Illumina reads and by Merqury (v1.3)(50) with 31-mer databases built from Illumina and HiFi data (singletons removed before merging). Completeness was assessed using Compleasm (v0.2.6)(26), which calculates the proportion of complete and single-copy BUSCOs to evaluate the presence of conserved orthologous genes. For sex chromosome evaluation, BUSCO genes specific to the opposite sex chromosome were excluded (i.e., Y-specific genes excluded for chromosome X evaluation, and X-specific genes excluded for chromosome Y evaluation).

Two additional DSAs we constructed–HCC1395BL (low contig N50) and HG008-N-D (assembly failure)–were of poor quality and therefore excluded from analysis. The remaining assemblies for COLO829BL, BL1437, BL2009,HCC1937BL, HCC1954BL and HG008-N-P, were used as diploid DSAs for somatic SV calling. For HG008, the DSA was built from HG008-N-P, but the matched normal sample for somatic SV calling was HG008-N-D because Illumina data for HG008-N-P were unavailable. Throughout the paper, these DSAs are referred to as COLO829, H1437, H2009, HCC1937, HCC1954, and HG008.

### Read mapping and variant calling

Short reads were aligned to GRCh38, CHM13, DSA Hap1 and DSA Hap2 with BWA-MEM (v0.7.17) (31), and PacBio HiFi and ONT reads were aligned with minimap2 (v2.28) (28). Long-read alignment were phased and haplotagged with Clair3 (v1.0.4)(51) and WhatsHap (v2.2)(52) following the Severus recommendation (53).

Somatic SVs were called with fifteen caller-platform combinations spanning three sequencing technologies and nine distinct tools (Figure 4a): DELLY v1.2.9(54), SvABA v1.1.3(55) and GRIDSS2 v2.13.2(56) (Illumina); Severus v1.2 (12), SAVANA v1.2.5 (57), Sniffles2 v2.4 (58) and nanomonsv v0.7.2 (59) (HiFi and ONT); and pbsv v2.9.0 (60) plus Sawfish v0.12.4 (61) (HiFi only). Each caller used matched tumor and normal BAMs as input and ran with default parameters. SVs from different call sets were considered equivalent when both breakpoints lay within a 500 bp distance (agnostic to SV type), a relaxed threshold that reduces representation bias across tools.

To construct a high-confidence set for each reference and sample, call sets were merged per haplotype and an SV was retained if supported by at least two of three sequencing platforms and by at least four of the fifteen calling strategies. This filtering approach balances sensitivity and specificity (12). The identical haplotype-specific pipeline was applied to GRCh38, CHM13, and two DSA haplotypes, ensuring that differences in SV discovery arise from reference choice rather than analytical artifacts. Finally, the two DSA haplotype-based high-confidence call sets were merged into a unified DSA-based set using EchoSV.

### EchoSV

Comparing SV call sets across references is non-trivial because the same variant can be represented differently on different genomes, with variations in contigs and coordinates of the breakpoints, SV type and length. While breakpoint coordinates can often be lifted-over in many regions, they fail in DRRs (differential reference regions) where sequences do not align uniquely between references. Local sequence differences can also cause the same event to be reported with different SV sizes or even types (Figure 6b). To address these challenges, we developed EchoSV, which captures how SVs “echo” across references through a hybrid workflow combining lift-over and graph-based matching. EchoSV accepts as input two or more SV call sets from the same sample and each generated on a different reference, and outputs either (1) a merged call set (e.g., consolidating the two DSA haplotype-based call sets into a unified set; Figure 1b) or (2) a comparison result (e.g., identifying overlapping and reference-exclusive SVs among GRCh38, CHM13, and DSA)

#### Classifying supporting reads across multiple platforms

Although a variant’s contig, coordinate, type and length may change from one reference to another, the reads that support this alternative allele remain the same(17). EchoSV therefore begins by collecting the complete set of read names that support each high-confidence SV across all available platform data from the tumor sample (to minimize coverage bias across platforms). We extracted reads with MAPQ*≥*1 whose SV signal shows breakpoints that lie within 500 bp of the given SV. Since most long or short reads span only a single SV, this union set of read names provides a reliable indicator of the SV’s identity, forming the basis for EchoSV’s accuracy.

#### Weighted complete bipartite graph

For every SV pair drawn from two different references, we quantified their similarity with an echo score, defined as the Jaccard index of their multi-platform supporting-read sets (i.e, the ratio of the intersection size to the union size). The score ranges from 0 (no shared reads, indicating unrelated events) to 1 (identical read support, strongly suggesting the same event). Using these scores, EchoSV constructs a complete weighted bipartite graph where each node represents an SV and every possible cross-reference SV pair is connected by an edge weighted by the echo score. This graph captures SV similarity with high fidelity and allows EchoSV to apply a globally optimal matching algorithm to identify the complete set of concordant SVs.

#### Finding concordant SVs

For SVs in regions that map uniquely between references (non-DRRs), breakpoint coordinates are lifted over and compared with a 500 bp tolerance. Pairs that match and have a non-zero echo score are immediately recorded as concordant and are excluded from the weighted bipartite graph. For the remaining SV pairs, EchoSV applies a maximum matching algorithm that selects a set of SV pairs that: (1) are non-overlapping, (2) each pair has an echo score > 0.5 (default `–-min-echo-score=0.5`), and (3) the total echo score is maximized across the graph. These selected pairs are added to the concordant set, yielding a robust and comprehensive match (robustness to the choice of `–-min-echo-score` is shown in Supplementary Table S4).

#### Haplotype-specific filtering for DSA consolidation

SVs detected on only one DSA haplotype may represent either true somatic events in haplotype-specific sequences (e.g., chromosome Y in male donors) or technical artifacts. To distinguish between these cases, EchoSV applies an additional filter that leverages the classified supporting reads. An SV exclusive to one haplotype is discarded as germline or mapping noise if the matched-normal alignments contain clear support (at least three supporting reads in both HiFi and ONT data). The retained variants are then merged into the unified DSA-based set.

#### Multi-reference comparison

EchoSV scales naturally to three or more references by chaining pairwise comparisons; we verified that the resulting concordant and reference-exclusive sets are effectively unchanged by the order in which the references are processed.

### SV annotation with repeat elements and overlapping gene

DSA haplotypes were annotated with RepeatMasker (v4.1.8)(30). An SV was flagged as repeat-associated when its breakpoints intersected any annotated repeat elements. To assign gene overlaps, RefSeq transcript sequences (excluding pseudogenes) were mapped to DSAs with BLAT (v35)(62), and SVs intersecting those alignments were recorded as gene-overlapping events.

### SV validation with assemblies

To evaluate each reference-exclusive SV, we assessed whether it’s supported by genome assemblies through aligning the assemblies to the corresponding reference. Tumor genomes were assembled using only PacBio HiFi reads with hifiasm (ploidy=2) (36) (as scaffolding using ONT provided little value for SV validation). As shown in Figure 5a, we check each SV with alignment of tumor genome assembly and then DSA. A reference-exclusive SV was classified as a true positive if it was reproduced in the tumor assembly alignment. If absent from the tumor assembly, we examined the DSA-to-reference alignments and determined that those variants present in the DSA were labeled as germline artifacts, while non-insertion variants co-localizing with breakpoints of large germline insertions were considered mapping artifacts. Variants not validated as either true or false positives were labeled unknown. Representative examples of true and false positives are shown in Supplementary Figure S11.

## Data availability

All raw sequencing data for COLO829, H1437, H2009, HCC1395, HCC1937, and HCC1954 are available through NCBI BioProject PRJNA1086849 (12); PacBio Revio HiFi reads for COLO829 and HCC1395 are in the PacBio public repository (44). HG008 tumor-normal trio data are from GIAB (45). Protein-coding gene-level TPM values for tumor cell lines were downloaded from the DepMap 24Q4 release (46), and the matched normal-tissue medians were taken from GTEx v10 (47).

## Code availability

EchoSV is publicly available at https://github.com/parklab/EchoSV.

## Acknowledgements

We thank Simon Chu for assistance with manuscript proofreading and Shannon Ehmsen for feedback on figures. This work was supported by grants from the US National Institutes of Health (UM1DA058230 and R01HG012573).

## Author contributions

Y.Z., H.L. and P.J.P. conceived the study. P.J.P. supervised the study. Y.Z. implemented computational methods and performed data analysis. H.Q. generated DSAs and evaluated their quality for the main analyses. Q.Q. generated DSAs using Verkko for the replication analysis. Y.Z. and P.J.P. drafted the manuscript with input from all authors. All authors read and approved the final paper.

