## Supplementary Figures for "Diploid donor-specific assembly enhances somatic structural variant detection in cancer genomes"

### Supplementary Figures for Zhang et al.

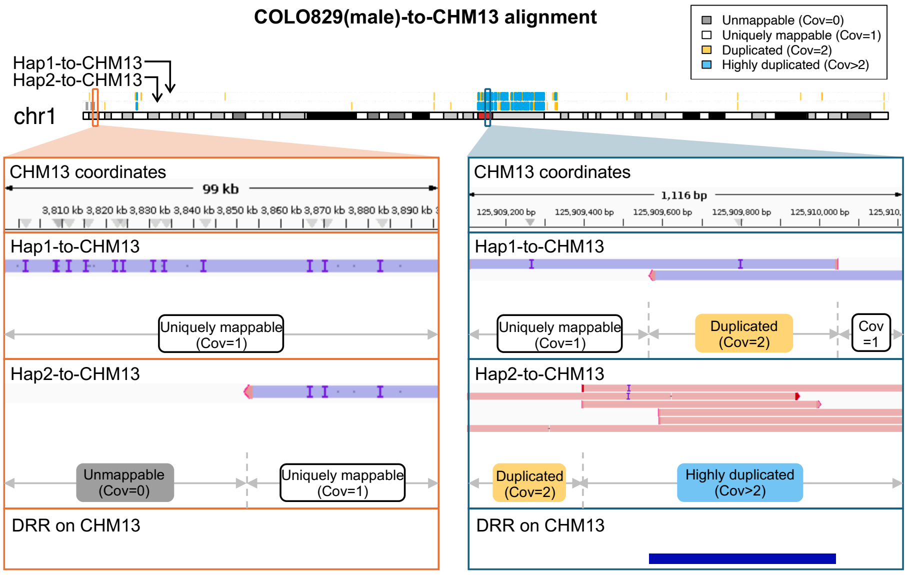


**Figure S1. Representative examples of CHM13 regions compared with DSA.**
Examples of **(left)** uniquely mappable (Cov=1) and unmappable (Cov=0) regions, and **(right)** duplicated (Cov=2) and highly duplicated (Cov>2) regions from DSA-to-CHM13 alignments. DSA contigs are shown in pink (forward strand) and purple (reverse strand) when aligned to CHM13. The final panel highlights DRRs (differential reference regions), defined as regions not uniquely mappable by either DSA haplotype.

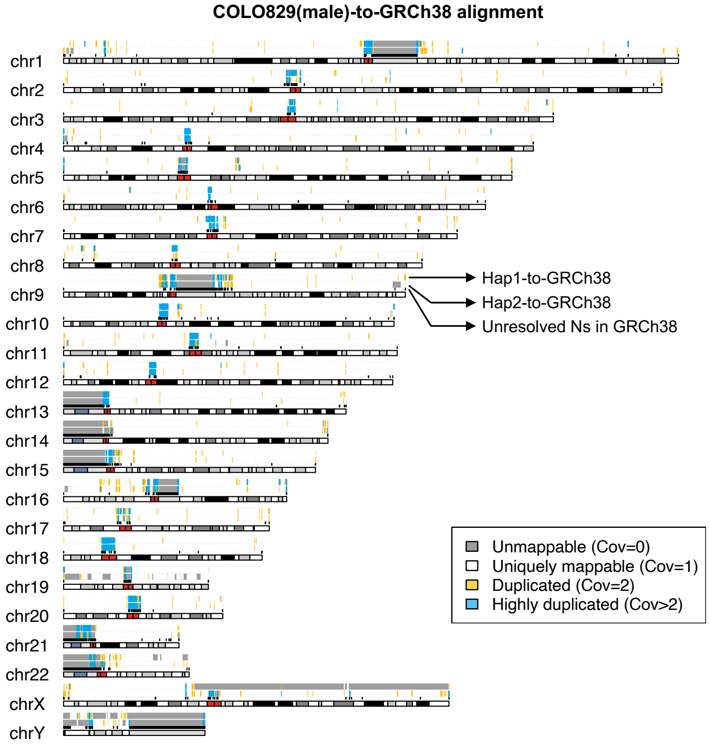


**Figure S2. Sequence comparison of COLO829 DSA with GRCh38.**
Assembly-to-assembly alignment of both haplotypes of the COLO829 DSA (male donor) against GRCh38 using minimap2, visualized with KaryoploteR (primary chromosomes only). Above each cytoband, two colored tracks show the four region classes (unmappable, uniquely mappable, duplicated, highly duplicated) for each DSA haplotype; a third black track highlights unresolved regions (Ns) in GRCh38.

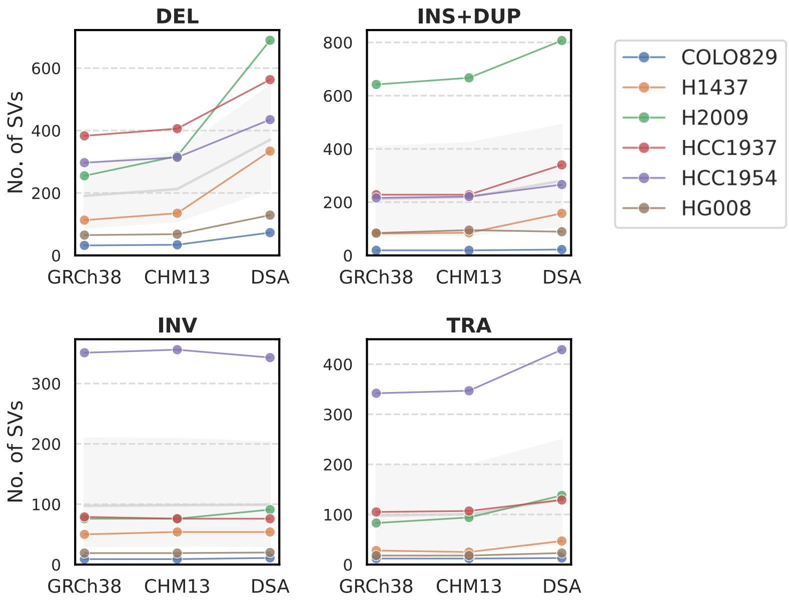


**Figure S3. Comparison of high-confidence SV counts across GRCh38, CHM13, and DSA by SV type.**
High-confidence SV counts on GRCh38, CHM13, and the consolidated DSA set, stratified by SV type. Grey lines and shaded areas show mean ± SD across six tumors; colors indicate the six tumor cell lines (legend, top right). Abbreviations: DEL, deletion; INS, insertion; DUP, duplication; INV, inversion; T, translocation.

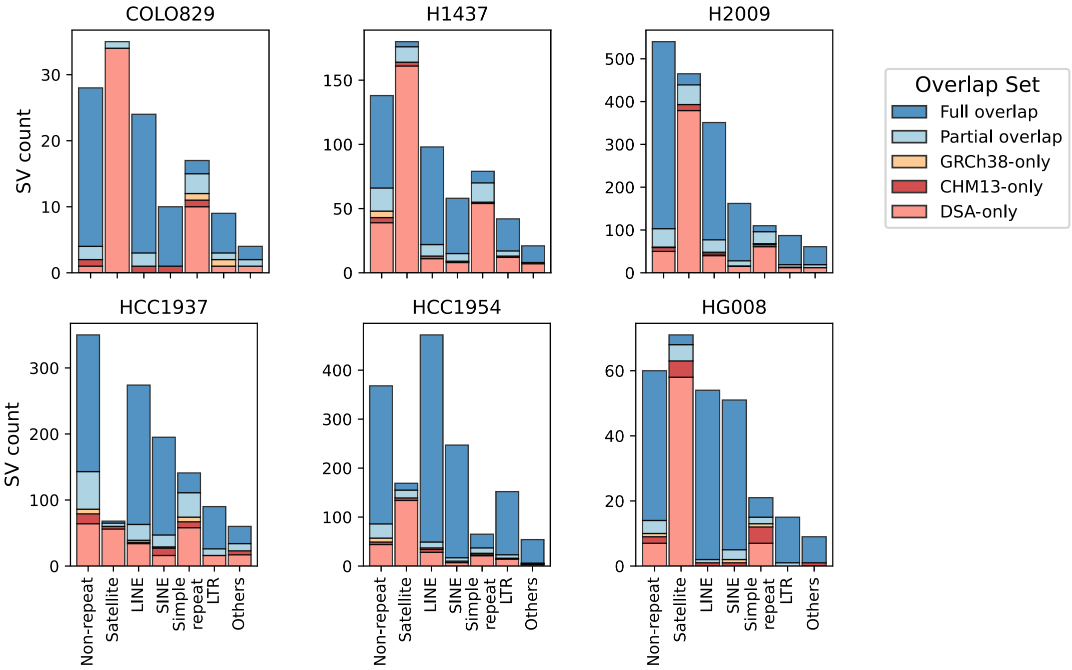


**Figure S4. SV counts by RepeatMasker class in each cell line.**
Counts of across-reference shared and reference-exclusive SVs in six cell lines, stratified by RepeatMasker class (extended from **Figure 4e**).

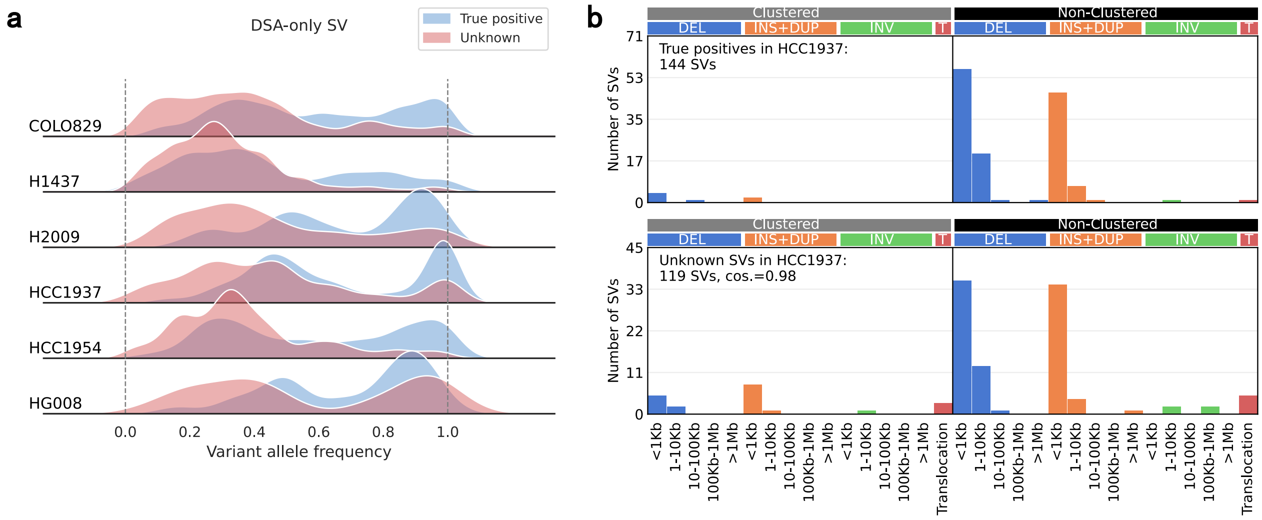


**Figure S5. VAF distribution and SV signature analysis of DSA-only SVs validated versus unknown.**
**a.** VAF distributions of tumor-assembly-validated versus unknown (unvalidated) DSA-only SVs across six tumor cell lines.
**b.** SV signatures in HCC1937 for validated versus unknown DSA-only SVs, stratified by clustering status (within 1 kb of another SV), SV type, and size. Abbreviations: DEL, deletion; INS, insertion; DUP, duplication; INV, inversion; TRA, translocation. The cosine similarity (cos.) between the two signatures of validated and unknown SVs is 0.98 (shown in the unknown panel); values for the other five cell lines were all >0.9.

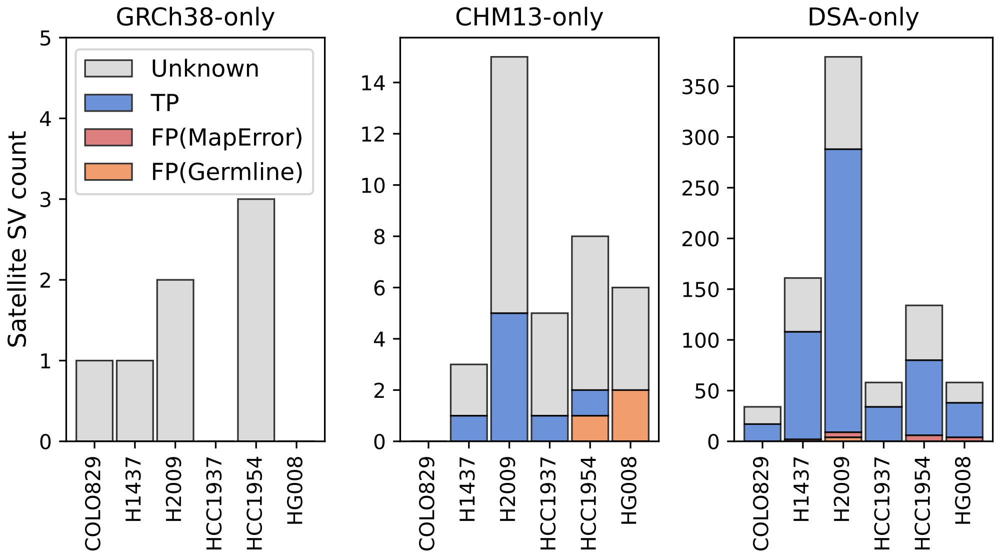


**Figure S6. Validation of reference-exclusive SVs in satellite regions.**
Stacked bar plots showing counts of reference-exclusive SVs by validation outcome. In satellite regions, 66% of DSA-only SVs were validated as true positives, compared with none for GRCh38-only SVs and 22% (roughly 1 per sample) for CHM13-only SVs. Exact counts are provided in Supplementary Table S7.

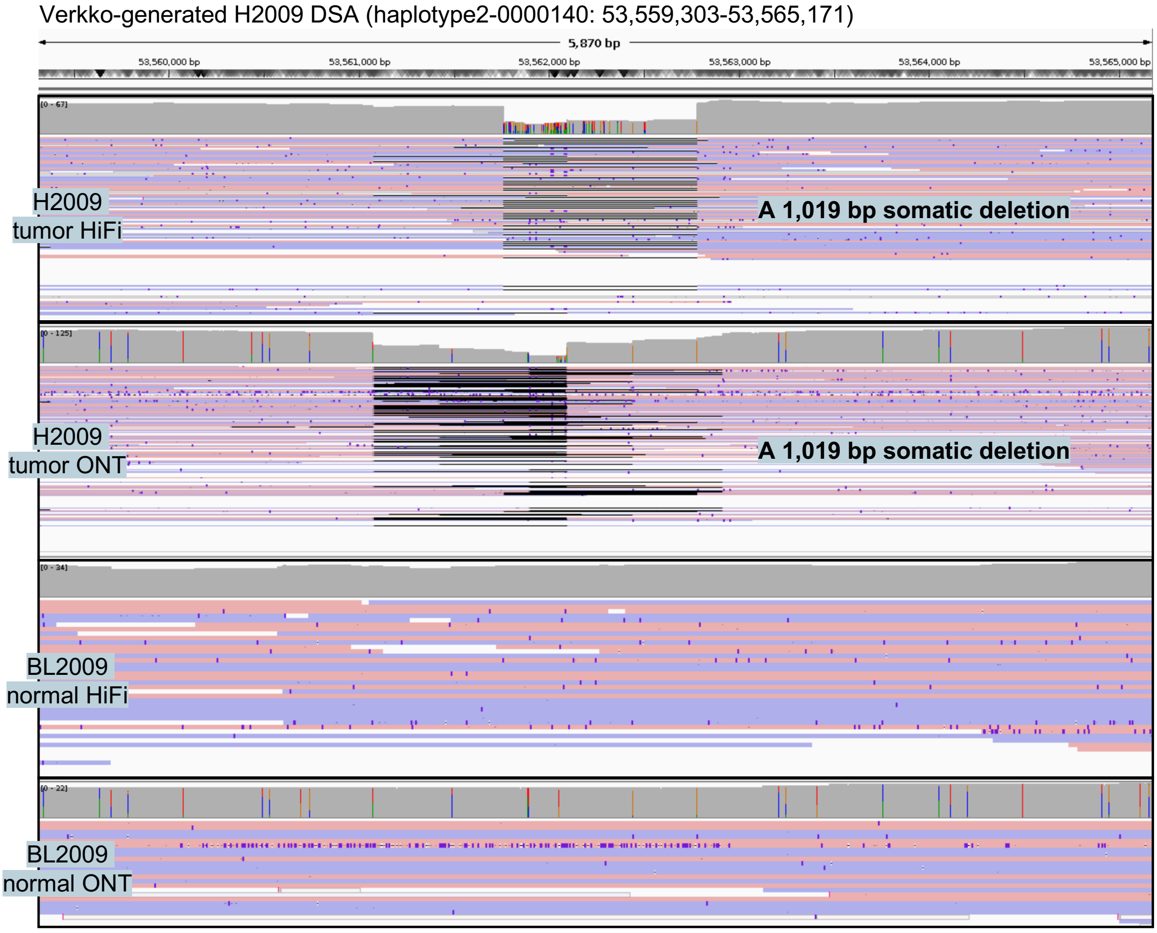


**Figure S7. Technical validation of the Figure 5d case using an alternative assembly and alignment pipeline.** This figure validates the 1,019 bp somatic deletion in a satellite repeat shown in **Figure 5d**. The SV was originally identified using a DSA created with hifiasm and alignments from minimap2. Here, we successfully reproduce the finding using an entirely independent pipeline: we generated a new DSA with Verkko (using HiFi, ONT, and additional Pore-C data) and mapped long reads with Winnowmap. This confirms that the detection of this SV is robust and not dependent on a specific choice of software. Breakpoints not exactly aligned in tumor read alignments are expected due to the inherent challenges of read alignment within satellite repeats. The four panels show IGV snapshots of the Winnowmap alignments for both HiFi and ONT reads against the diploid Verkko DSA (including both haplotypes and unassigned contigs). Reads are colored by strand (pink: forward, purple: reverse), with white indicating ambiguously mapped reads (MAPQ=0).

Verkko-generated H2009 DSA (haplotype1-0000026:3,893,426-3,905,727)

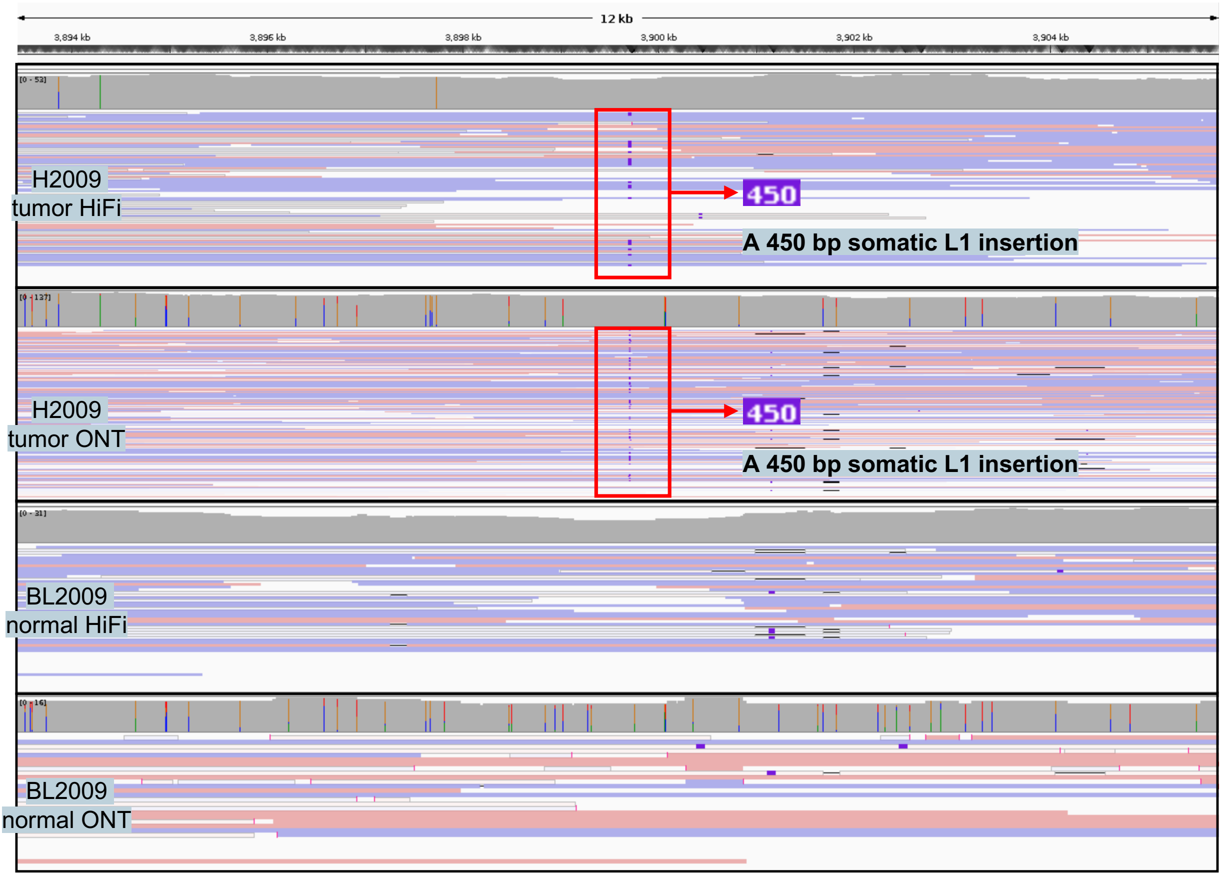


**Figure S8. Technical validation of the Figure 6c case using an alternative assembly and alignment pipeline.** This figure validates the 450 bp somatic L1 insertion in a satellite repeat shown in **Figure 6c**. The SV was originally identified using a DSA created with hifiasm and alignments from minimap2. Here, we successfully reproduce the finding using an entirely independent pipeline: we generated a new DSA with Verkko (using HiFi, ONT, and additional Pore-C data) and mapped long reads with Winnowmap. This confirms that the detection of this SV is robust and not dependent on a specific choice of software. The four panels show IGV snapshots of the Winnowmap alignments for both HiFi and ONT reads against the diploid Verkko DSA (including both haplotypes and unassigned contigs). Reads are colored by strand (pink: forward, purple: reverse), with white indicating ambiguously mapped reads (MAPQ=0).

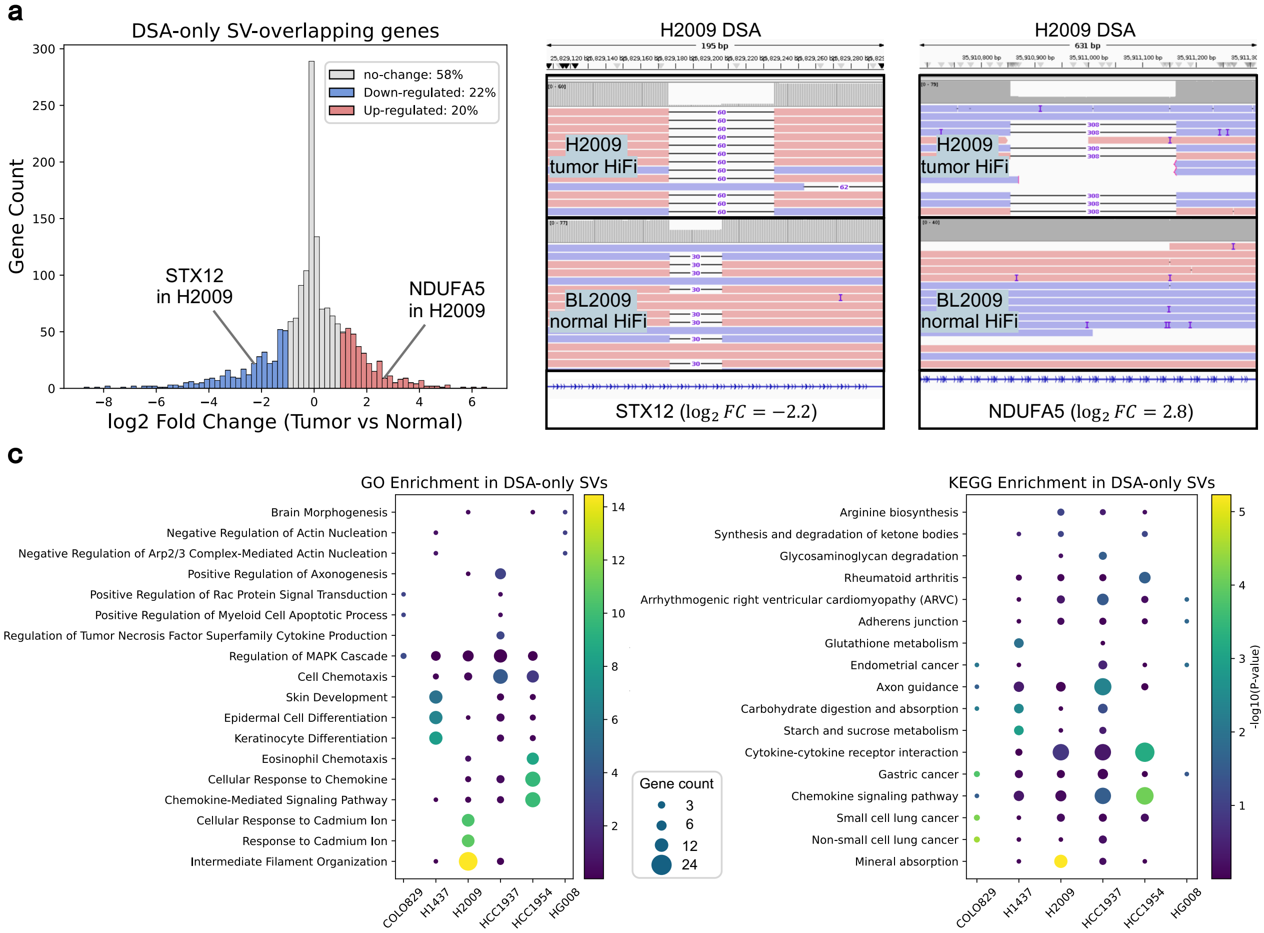


**Figure S9. Functional annotation of genes overlapping DSA-only SVs.**

**a.** Distribution of protein-coding genes intersecting DSA-only SVs by tumor–normal expression change ($\log_2$FC). Up-regulated genes ($\log_2$FC > 1) are shown in red, down-regulated genes ($\log_2$FC < –1) in blue, and unchanged genes ($|\log_2$FC$| < 1$) in grey. IGV examples include STX12, which overlaps a 60 bp DSA-only deletion in H2009 and is downregulated in the tumor, and NDUFA5, which overlaps a 308 bp DSA-only deletion in H2009 and is upregulated.

**b.** GO and KEGG pathway enrichment of genes intersecting DSA-only SVs. For each cell line, the top three enriched terms (ranked by $p$-value) are marked, with dot size indicating the number of overlapping genes (size legend in the center) within that category, and color intensity reflects significance.

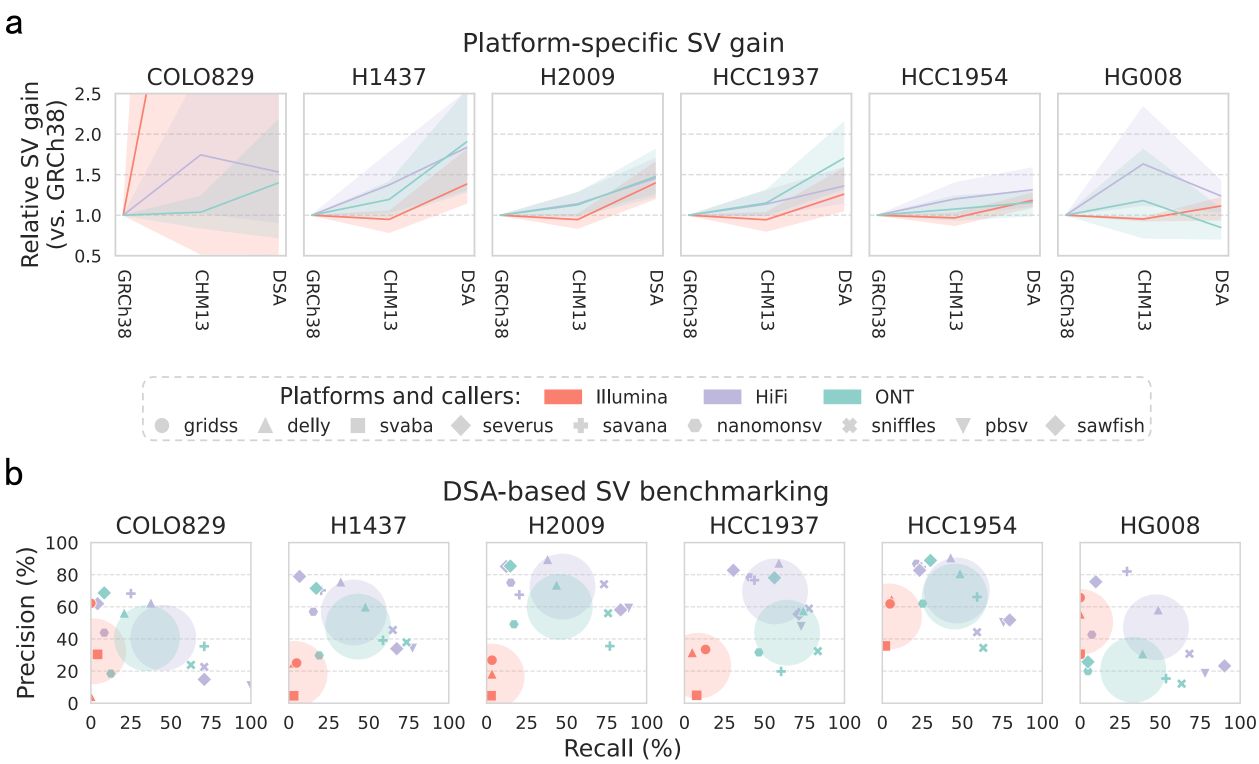


**Figure S10. Long-read somatic SV calling benefits more from using DSAs as reference than short reads.**

**a.** Mean SV gain by platform, normalized to GRCh38 (set to 1). Lines indicate the average count across callers within each platform (three Illumina, seven HiFi, five ONT); shaded areas show mean±SD.

**d.** Precision–recall analysis of 15 calling strategies on the DSA benchmark. Colors indicate sequencing platforms; markers represent individual callers, and large circles denote platform means. Recall was calculated against 832 tumor-assembly-confirmed DSA-only SVs, reflecting each caller's ability to detect true DSA-specific events. Precision was estimated relative to the DSA-based high-confidence set, indicating the extent of platform- and caller-specific artifacts.

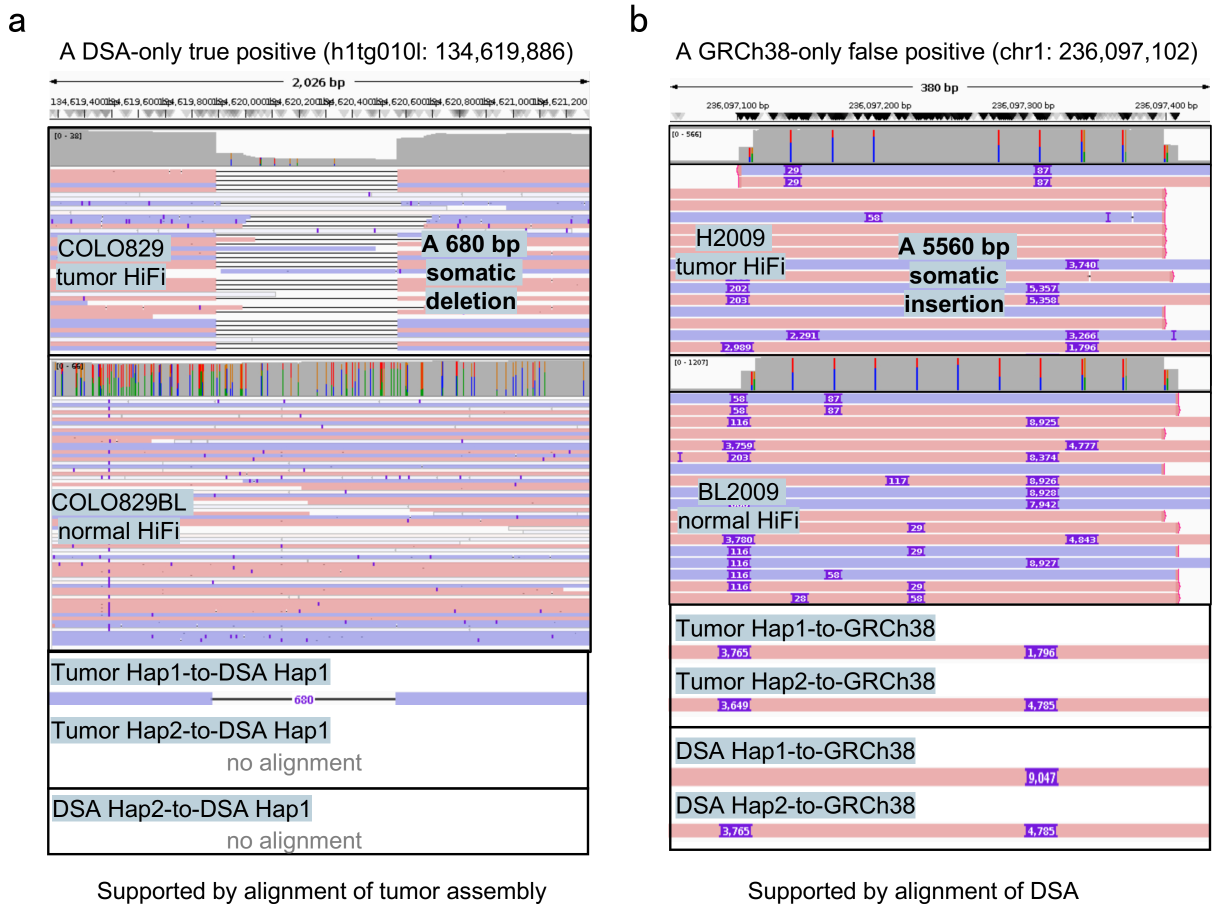


**Figure S11. Examples of reference-exclusive SVs validated as true or false positives, visualized in IGV.**
**a.** A 680 bp DSA-only somatic deletion validated as a true positive, as it is reproduced in the tumor genome assembly.
**b.** A 5,560 bp GRCh38-only somatic insertion validated as a false positive, as it is present in the DSA.
Both examples are high-confidence SVs, supported by at least two sequencing platforms and four calling strategies. For each case, HiFi tumor and normal read alignments, together with alignments of the tumor assembly and DSA, are shown. Read and contig alignments are colored pink (forward strand) and purple (reverse strand).
